# Leveraging neural networks to correct FoldX free energy estimates

**DOI:** 10.1101/2024.09.23.614615

**Authors:** Jonathan E. Barnes, L. América Chi, F. Marty Ytreberg, Jagdish Suresh Patel

## Abstract

Proteins play a pivotal role in many biological processes, and changes in their amino acid sequences can lead to dysfunction and disease. These changes can affect protein folding or interaction with other biomolecules, such as preventing antibodies from inhibiting a viral infection or causing proteins to misfold. The ability to predict the effects of mutations in proteins is crucial. Although experimental techniques can accurately quantify the effect of mutations on protein folding free energies and protein-protein binding free energies, they are often time-consuming and costly. By contrast, computational techniques offer fast and cost-effective alternatives for estimating free energies, but they typically suffer from lower accuracy. Enhancing the accuracy of computational predictions is therefore of high importance, with the potential to greatly impact fields ranging from drug design to understanding disease mechanisms. One such widely used computational method, FoldX, is capable of rapidly predicting the relative folding stability (ΔΔG_fold_) for a protein as well as the relative binding affinity (ΔΔG_bind_) between proteins using a single protein structure as input. However, it can suffer from low accuracy, especially for antibody-antigen systems. In this work, we trained a neural network on FoldX output to enhance its prediction accuracy. We first performed FoldX calculations on the largest datasets available for mutations that affect binding (SKEMPIv2) and folding (ProTherm4) with experimentally measured ΔΔG. Features were then extracted from the FoldX output files including its prediction for ΔΔG. We then developed and optimized a neural network framework to predict the difference between FoldX’s estimated ΔΔG and the experimental data, creating a model capable of producing a correction factor. Our approach showed significant improvements in Pearson correlation performance. For single mutations affecting folding, the correlation improved from a baseline of 0.3 to 0.66. In terms of binding, performance increased from 0.37 to 0.61 for single mutations and from 0.52 to 0.81 for double mutations. For epistasis, the correlation for binding affinity (both singles and doubles) improved from 0.19 to 0.59. Our results also indicated that models trained on double mutations enhanced accuracy when predicting higher-order mutations (such as triple or quadruple mutations), whereas models trained on singles did not. This suggests that interaction energy and epistasis effects present in the FoldX output are not fully utilized by FoldX itself. Once trained, these models add minimal computational time but provide a substantial increase in performance, especially for higher-order mutations and epistasis. This makes them a valuable addition to any free energy prediction pipeline using FoldX. Furthermore, we believe this technique can be further optimized and tested for predicting antibody escape, aiding in the efficient development of watch lists.

## Introduction

Proteins are essential to many biological processes, from structural support to cellular interactions. Their diverse functions are determined by their unique amino acid sequences and three-dimensional structures. Mutations in these sequences, whether single or multiple, can significantly impact biophysical properties like folding stability and binding affinity, thereby altering protein function. Moreover, the effects of multiple mutations may not be additive, leading to complex epistasic effects. Protein mutations are implicated in several diseases, including prion diseases, genetic disorders, cancer, and viral infections.^1–6^ Viruses, in particular, can mutate their surface proteins to enhance their ability to infect hosts or evade treatments like antibodies.^7–10^ Conversely, a mutation might impair proper protein folding, negating any binding affinity advantages.^11^ We can also harness mutations to improve therapeutics, such as enhancing antibody binding affinity.^12^ Given these factors, it is crucial to predict the effects of mutations on protein biophysical properties quickly and accurately to advance human health.

Several experimental techniques, such as surface plasmon resonance and isothermal titration calorimetry, are commonly employed to evaluate the impact of mutations on binding affinity.^13^ Additionally, methods like differential scanning calorimetry, circular dichroism, and UV absorbance are utilized to assess the folding stability of biomolecules.^14^ While these techniques provide reliable data, they are limited by their low throughput and high operational costs.

To address these limitations, various computational methods have been developed that offer a broader range of capabilities. The current state-of-the-art computational approaches for free energy calculations are typically categorized into two groups: ”slow” and ”fast” methods. Slow methods, such as the Molecular Mechanics Poisson-Boltzmann Surface Area (MMPBSA) approach, are known for their high accuracy, with reported correlation coefficients up to 0.92 when compared to experimental data for protein-ligand binding interactions.^15^ However, these methods are constrained by their computational intensity and low throughput.

In contrast, fast methods employ semi-empirical, statistical, or physics-based algorithms, including advanced machine learning-based approaches.^16–25^ These fast methods, while generally less accurate, provide significantly higher throughput, making them more suitable for large-scale applications, such as screening thousands of potential mutations. Consequently, for high-throughput requirements, fast methods are often preferred due to their balance between speed and computational efficiency

Furthermore, both fast and slow methods are rarely developed for or tested against more than single point mutations. This is due to multiple mutations potentially involving epistasis, making prediction more difficult as epistasis is still not fully understood. Wells in 1990^26^ made the earliest observations of potential explanations like separation distance, conformational perturbations, and charge. We recently used statistical methods to try and discover mechanistic insights.^27^ Whereas others have used machine learning to attempt to classify pairs of mutations as epistatic.^28^ As it stands, few of the aforementioned methods are rigorously tested for multiple mutations. Some, like DDAffinity from Yu et al.^17^ or mmCSM-PPI by Rodrigues et al.^29^ are tested and evaluated for their performance on multiple mutations, though these tend to show minimal performance improvement on larger broad datasets. These methods are also generally evaluated on multiples as a whole, not broken down by order.

Among the “fast methods”, FoldX^30^ and FoldX-based approaches^31^ are widely used for predicting the effects of mutations on free energies. ^32^ FoldX is a software package that relies on a semi-empirical force field and multiple energy terms to predict free energies.^33^ In 2020, Gonzalez et al. evaluated 8 different methods for predicting protein-protein binding affinity for single mutations, including FoldX and FoldX-based approaches.^34^ They discovered that the best method depends on the problem at hand, with the peak predictive accuracy across all methods being 0.49. Some of the most accurate results for antibody-antigen systems were obtained by combining molecular dynamics with FoldX, with a correlation coefficient (r) of 0.39. However, this method was also the most computationally expensive. Overall, these results suggested that while these methods can quickly and accurately predict stabilizing versus destabilizing mutations, they are less precise in predicting actual binding affinities, indicating room for improvement.

In this study, we have developed an approach that combines energy terms (referred as features) generated by the FoldX program and a neural network to improve predictions of protein-protein binding affinity for both single and higher order mutations. This approach is simple, not requiring any external data or intense computation, and improves overall accuracy for single and multiple mutations.

## Methods

In this work we improved FoldX predictions using neural networks to estimate a correction factor. Our process first involved acquiring the largest datasets of experimental free energy of protein folding and binding available and the corresponding 3D crystal structures. We then used a custom FoldX protocol to predict the free energy of folding and binding for the available data, extracting the FoldX energy terms for each estimate. These free energy predictions and energy terms then went through a feature selection process, leaving us with final feature sets. Neural networks were then optimized and trained on these feature sets to predict correction factors for folding and binding, greatly improving the original FoldX estimates. Lastly, we explored our methods’ capability of predicting higher-order mutations and if models trained a lower orders of mutations had a predictive capability that transferred to higher orders.

To improve the accuracy of FoldX predictions, we developed an approach that combines energy terms (refered as features) generated by FoldX program and a neural network. Resulting models were systematically tested for both single and higher-order mutations. The detailed methodology is described below.

### Dataset Curation and Preprocessing

Our process first involved acquiring the largest datasets of experimental free energy of protein folding and binding available and the corresponding 3D crystal structures. Experimental binding affinity data was acquired from SKEMPIv2^35^ and folding data from ProTherm 4.0.^36^ Both datasets needed processing to be usable. For SKEMPIv2, after downloading the entire dataset as a CSV file, free energy values were derived from affinity values K*_D_* using the following relationship:

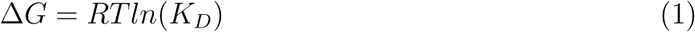

ProTherm data required cleaned and converted to a column-based format as it is provided as a text file with blocks of entries rather than in a tabulated form. After formatting, both SKEMPIv2 and ProTherm ΔG data for mutant and wildtype were combined to generate ΔΔG values for each system and mutation:

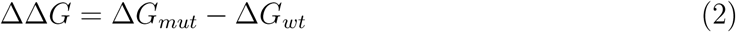

Where ΔG*_mut_* and ΔG*_wt_* are the free energies of the mutant and wildtype respectively. For folding stability, we compiled single, double, and an ”all multiple” mutation datasets. For binding affinity we compiled data for single, double, triple, and quadruple mutations in addition to an ”all multiples” dataset. The ”all multiples” datasets are comprised of double thru the highest order available. These full datasets are outlined in 1.

After curating all of our free energy datasets, 3D crystal structures were acquired from the RCSB Protein Databank^37^ for all systems. These crystal structures were then processed using PDB Fixer^38^ to add missing residues, heavy atoms, and correct any other crystallographic errors to then be used as inputs for FoldX and any other modeling programs.

Lastly, we also explored epistasis prediction, which requires data for higher-order mutations and their constituent lower-order mutations. For double mutations, epistasis is defined by:

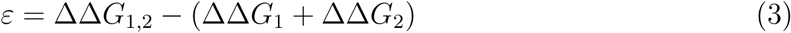

Where *ε* is epistasis, or the deviation in additivity, and ΔΔG_1,2_, ΔΔG_1_, ΔΔG_2_ are the free energy values for the double mutant, and constituent single mutations respectively. Since we have a large amount of binding affinity data, we used those datasets (and none from folding stability) for all epistasis analysis. We created a subset of double mutations, referred to as D558, consisting of all double mutations with the constituent singles available. We also created a separate single mutation dataset comprised of the corresponding constituent single mutations referred to as S487.

**Figure 1:**
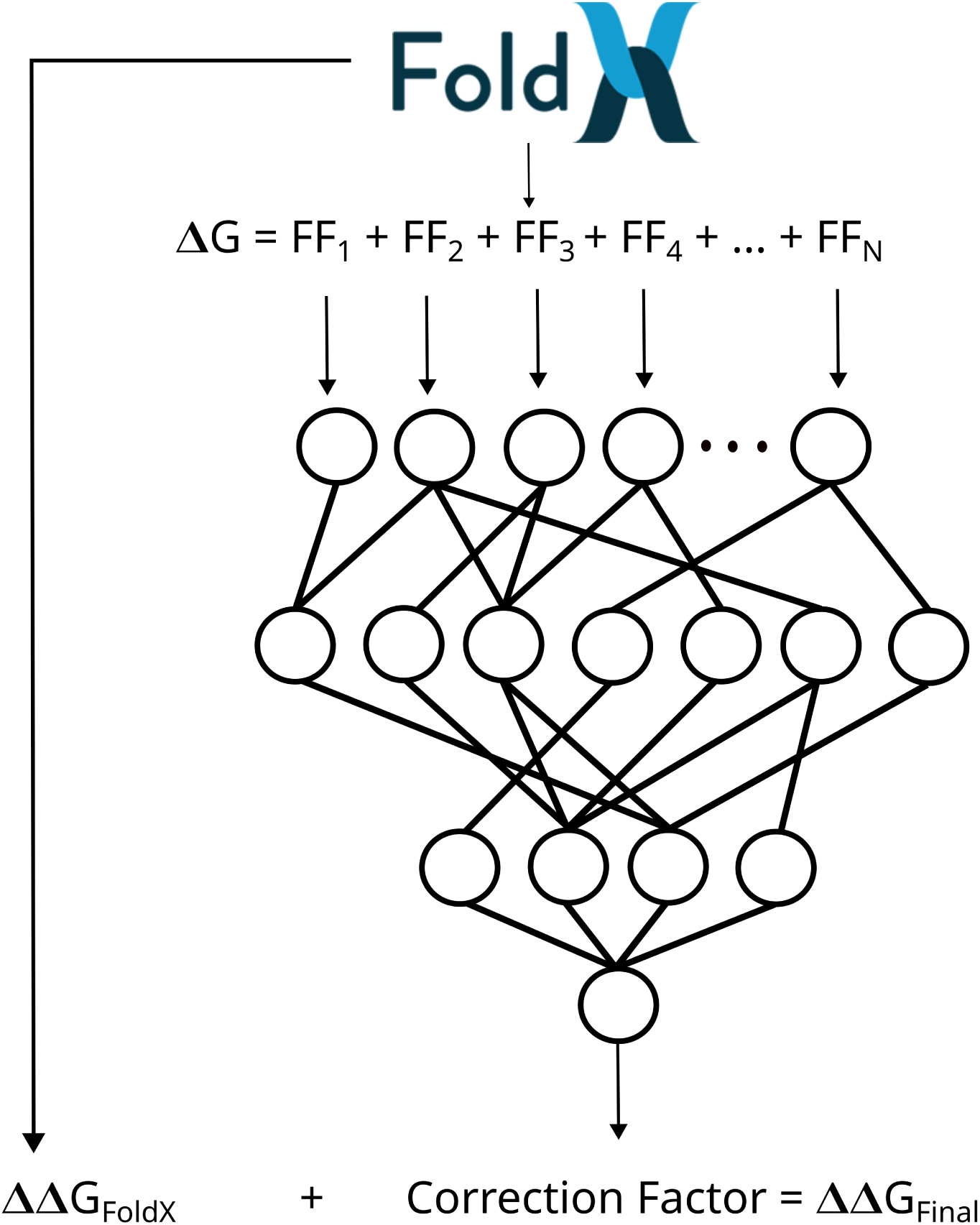
General methodology flowchart. FoldX is used to perform mutations, predict ΔΔG, and to provide the energy terms to feed into the corrected neural network.

### FoldX Calculations

To obtain predicted ΔΔG values and associated free energy terms, FoldX calculations were performed using FoldX v4.0^30^ and our previously published protocol. Prepared structures were minimized using *RepairPDB* six times in succession to ensure convergence. Mutations were then generated using *BuildModel*, to create both single, double, and all higher-order mutations for free energy prediction. For binding affinity calculations, *AnalyseComplex* was used on the non-mutated wildtype and mutated structures to calculate the ΔG*_wt_* and ΔG*_mut_* estimates and then ΔΔG*_bind_* values were calculated. Similarly, for folding, we used the *Stability* command to predict ΔG*_fold_* for wildtype and mutants, to then generate ΔΔG*_fold_* estimates. The resulting free energy predictions and energy terms were then compiled for feature selection.

### Feature Extraction and Selection

The features used in this study were derived exclusively from FoldX, without incorporating data from other computational methods or *a priori* information about the proteins or amino acids being evaluated. Specifically, the complete feature set included FoldX predictions for ΔG and ΔΔG, as well as the individual energy terms contributing to these predictions. We assessed each feature for variation and correlation with experimental free energy measurements. Features that showed no correlation and no variation across all datasets were excluded. For binding affinity, the excluded features were: loop entropy, disulfide, partial covalent interactions, and Entropy Complex. For folding stability, we excluded: water bonds, loop entropy, kn electrostatic, partial covalent interactions, and Entropy Complex. Additionally, we removed the G estimates. The final dataset contained 21 columns for binding and 20 for folding. These columns included the Protein/System ID, mutation(s), FoldX ΔΔG estimate, energy terms, experimental ΔΔG (reference), and the response variable. The response variable was defined as the difference between the experimental and FoldX-predicted ΔΔG values (a correction factor), as this approach yielded more accurate results and was more physically meaningful than predicting experimental ΔΔG directly.

### Machine Learning

We next utilized neural networks to build models capable of predicting the correction necessary for FoldX outputs. We chose to use neural networks as they provide a framework to learn complex relationships between features at a high level and do so unsupervised. Furthermore, our datasets are fairly simple and do not require extensive resources as all our models were trainable in reasonable time frames on CPUs rather than requiring GPU resources.

Keras TensorFlow v2.16.1 served as the primary framework for our neural networks. To determine the optimal network structure, we employed the Keras-Tuner Python module v1.4.7. The hyperparameters we tuned included the size of the hidden layer(s), activation function(s), dropout layer rate, learning rate parameters, and the weight decay parameter. Dropout layers and decaying weights were used as regularization to reduce overfitting.

We evaluated three network structures to identify the optimal configuration: one with a single hidden layer, another with two hidden layers and an intermediate dropout layer, and a third with an arbitrary number of layers (up to five) as a hyperparameter with dropout. We used AdamW as the optimizer with decaying weights. The learning rate was adjusted via exponential decay with a decay rate of 0.95 and 100,000 decay steps, with the initial learning rate determined by Keras-Tuner.

We used the mean absolute error (MAE) as the evaluation metric for loss, with a 10% validation split for training. The tuning process was conducted with a max trials setting of 500 and executions per trial set to 3, using RandomSearch as the tuner. Although BayesianOptimization was considered, it was ultimately not used due to its tendency to get trapped in local minima. The maximum number of epochs for each run was 5000, with an early stop callback function set to a patience of 50, monitoring validation loss.

Once the tuning process was complete, we evaluated the results and selected the final model from the top three performing sets of hyperparameters, considering both simplicity and performance. We also checked for overfitting by plotting the training versus validation loss. The production model was then trained using the optimized parameters, following the same test-train split, maximum number of epochs, and early stop monitoring as in the initial tuning phase. By employing this thorough and systematic approach, we ensured that the resulting model was both robust and efficient, with minimal risk of overfitting and optimal performance. Final models are outlined in Tables 2 3.

Cross validation was used to confirm the generalizability of the methodology and optimal structure. The optimized hyperparameters were used with randomized test-train splits 100 times, this provided a simulation of how our models perform trained and evaluated on different datasets. The resulting Pearson correlation and Spearman rank performance were then summarized and given as a distribution. Final models were then created using the entire corpus of available data to apply to future external datasets.

We further tested models and their capability of generalizing prediction improvement to different orders of mutations. The final model trained on singles was applied to doubles, triples, and quadruples. The final model for doubles was applied to singles, triples, and quadruples. A model was also trained on triple mutations and evaluated on all other mutation sets. A model was not trained on quadruples due to only having 255 mutations available. For folding, we only built a model on single mutations due to a significantly small corpus available for double mutations.

For epistasis prediction, we followed a separate validation-based approach. The constituent single mutation subset (S487) was treated as a test set for a single mutation model. For the double mutation dataset (D558) we performed the full model selection pipeline outlined above, then performed 100-fold cross validation, saving each prediction for each validation run. The validation test sets were then averaged and used as values for epistasis predictions. Averaging was chosen as it smooths out outliers but doesn’t fully discount their contributions.

## Results and discussion

To build corrective models for FoldX, we first curated existing datasets of mutations with experimentally measured protein folding (ProTherm4) and binding (SKEMPIv2) free energy values. We then used FoldX to predict the free energies of binding and folding for all mutations available. The resulting output was used as features to train deep learning models. We then performed a hyper-parameter search detailed in the Machine Learning section of the methods to arrive at the best parameters for each model shown in Table 1. These models were then used on the full datasets, in validation, and applied to epistasis calculations.

**Table 1:** Summary of datasets used.

| **SKEMPIv2 (binding)** |  | **ProTherm 4.0 (folding)** |  |
| --- | --- | --- | --- |
| **Number of Mutations** | **Number of Data** | **Number of Mutations** | **Number of Data** |
| All | 6399 | All | 2340 |
| Singles | 4588 | Singles | 1860 |
| All multiple | 1690 | All multiple | 478 |
| Doubles | 858 | Doubles | 309 |
| Triples | 325 |  |  |
| Quadruples | 254 |  |  |

As shown in Tables 2 and 3, for all cases the best models consisted of two layers. For folding the initial learning rate was much smaller than for either binding model at 0.0005. The weight decay was also double the binding model values. The layers for folding were also nearly identical in size with no dropout layer, potentially increasing the possibility of over-fitting even if the loss plot does not indicate the problem [supp]. For binding, both single and double models contained a dropout layer with a dropout rate of 30%. Both models had the same weight decay of 0.004 with the double mutation model having double the initial learning rate. One interesting observation is the network dimensions are almost inverted between the single and double models, that is the singles model had a larger first layer whereas the double model had a larger second layer.

**Table 2:** Single mutation model hyperparmeters after RandomSearch process. Models consisted of two layers, including a drop-out layer.

| **Folding Single Mutations** | 1763 datapoints | **Binding Single Mutations** | 3855 datapoints |
| --- | --- | --- | --- |
| Random Seed: | 42 | Random Seed: | 42 |
| Train Size: | 1410 | Train Size: | 3084 |
| Test Size: | 353 | Test Size: | 771 |
| **Model:** |  | **Model:** |  |
| Initial Learning Rate | 0.0005 | Initial Learning Rate | 0.001 |
| Weight Decay | 0.008 | Weight Decay | 0.004 |
| **Layer 1 Activation** | **Layer 1 size** | **Layer 1 Activation** | **Layer 1 size** |
| gelu | 496 | swish | 432 |
| **Dropout** | 0 | **Dropout** | 0.3 |
| **Layer 2 Activation** | **Layer 2 size** | **Layer 2 Activation** | **Layer 2 size** |
| elu | 512 | softplus | 144 |

**Table 3:** Double mutation model hyperparmeters after RandomSearch process.

| **Binding Double Mutations** | 814 Datapoints |
| --- | --- |
| Random Seed: | 42 |
| Train Size: | 651 |
| Test Size: | 163 |
| **Model:** |  |
| Initial Learning Rate | 0.002 |
| Weight Decay | 0.004 |
| **Layer 1 Activation** | **Layer 1 size** |
| sigmoid | 272 |
| **Dropout** | 0.3 |
| **Layer 2 Activation** | **Layer 2 size** |
| tanh | 416 |

Table 4 summarizes our model performance for the main datasets tested. Second and third columns in the table shows similarity in FoldX performance observed between the full dataset and the test sets indicates that our test subsets are fairly representative of the full dataset in terms of distribution. For single mutation models, we see significant performance increases with the Pearson correlation more than doubling for folding and a greater than 60% improvement for binding affinity. Most of the observed corrections can be attributed to outliers being corrected [supp]. We also observe some significant outliers in FoldX predictions, such as one or two mutations having an estimated ΔΔG of +/- 20 kcal/mol, likely driving part of the poor linear correlation. Since we had a large enough corpus available, we also explored higher-order mutations in the case of binding affinity. For double mutations, we see a 60% improvement to FoldX on its own. We also tested a generalized multiple mutation model, consisting of all higher-order mutations in one dataset not divided by order. In this case, we see a similar performance boost to double mutations.

**Table 4:** Table of primary results for model corrections. Results are given for the Pearson correlation and Spearman rank. Models were trained and evaluated on the same dataset (e.g. - binding single mutations was a model trained on single mutations and evaluated on binding single mutations)

| **Pearson Correlation** |  |  |  |
| --- | --- | --- | --- |
| **Dataset and Model** | **FoldX (full data)** | **FoldX (test set)** | **Model (test set)** |
| Folding single mutations | 0.36 | 0.30 | 0.66 |
| Binding single mutations | 0.39 | 0.37 | 0.61 |
| Binding double mutations | 0.53 | 0.52 | 0.81 |
| Binding all multiples | 0.52 | 0.49 | 0.80 |
| **Spearman Rank** |  |  |  |
| **Dataset and Model** | **FoldX (full data)** | **FoldX (test set)** | **Model (test set)** |
| Folding single mutations | 0.50 | 0.52 | 0.63 |
| Binding single mutations | 0.42 | 0.45 | 0.59 |
| Binding double mutations | 0.56 | 0.57 | 0.77 |
| Binding all multiples | 0.58 | 0.56 | 0.80 |

Figure 2 shows the resulting epistasis prediction results. Left is the scatter plot of the averaged model corrected *ε* values compared to the experiment and right shows the separate validation subsets Pearson correlation to each respective subsets experimental data. In spite of the compounded error for the double mutant and constituent single mutations, we see significant improvement in predicting epistasis with a near three-fold increase in Pearson performance. FoldX, as previously mentioned, tends to predict additivity. This is evidenced by its values making almost a flat line near zero. Epistasis as an effect is still not fully understood. The top contributors are believed to be separation distance, charge interactions, and conformational perturbations.^26^ If we consider the terms being incorporated in our models the best case could be made for charge interactions.

**Figure 2:**
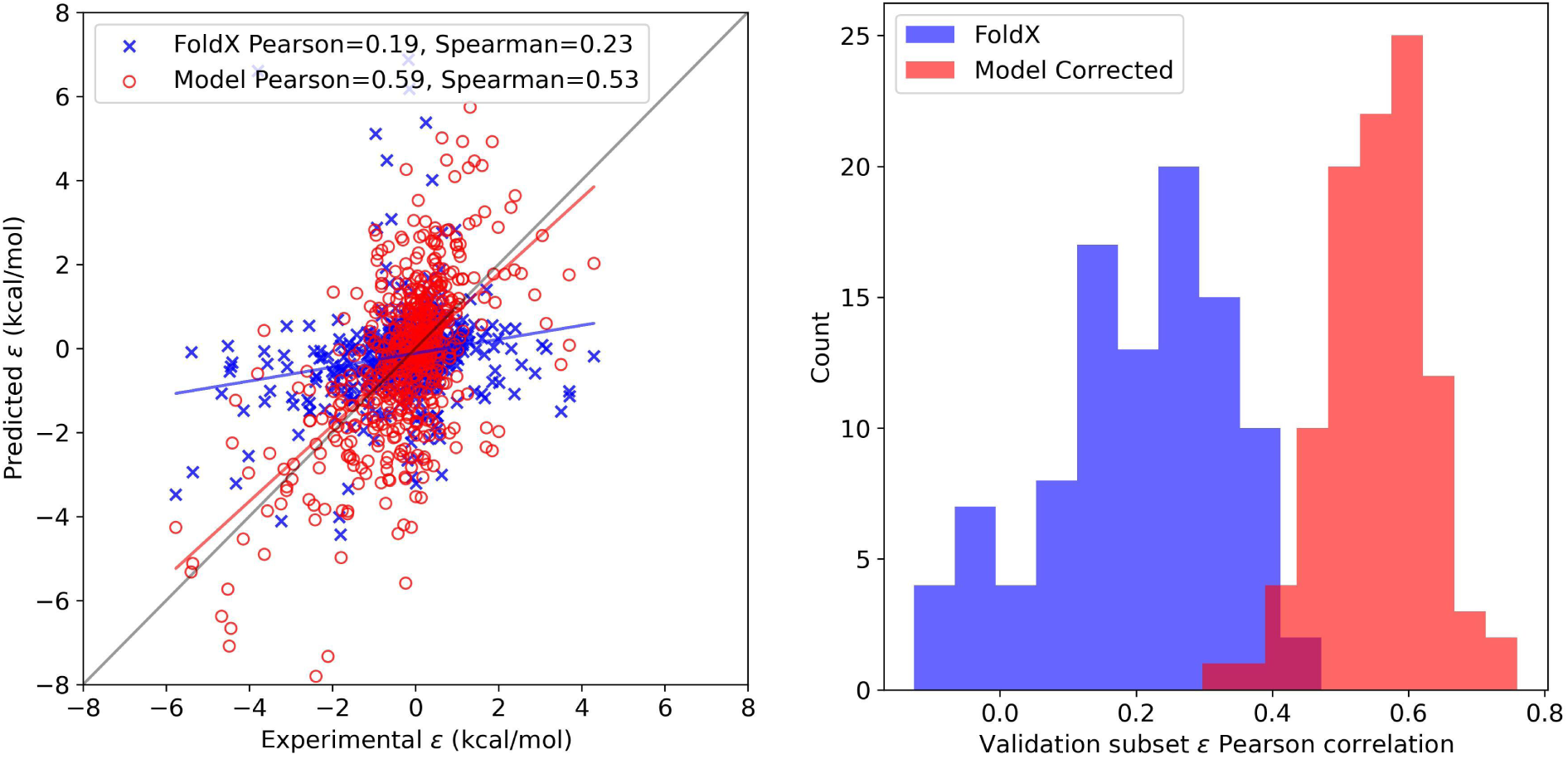
Results of epistasis prediction. Scatter plot (left) shows how well our single and double mutation models are capable of correcting epistasis. These results are comprised of test set predictions for 100 validation runs. The individual performance for each test set is shown on the right.

Figure 3 shows the model performance graphically for the subset trained on and applied to the other three datasets tested with the largest corpus available. The main observation is that the trained models perform better than plain FoldX, in all cases with an improvement of 0.2 or better for the Pearson correlation. In the case of singles, we see some small corrections to outliers but overall not a significant visual improvement. When we look at higher-order mutations the single mutation model performs marginally better but without a significant improvement. This is repeated for triples and quadruples where while there is usually an improvement – barring the case of triples where performance went down – the improvements observed are within error margins. For double mutations, we see a more significant improvement (Pearson=0.81 vs 0.52). Interestingly, in the case of the doubles model is that we see negligible improvement for the singles mutation dataset, however in the case of triples we see a decent performance boost (Pearson=0.81 vs 0.65). Lastly, for quadruples, we see very minimal improvement with the Spearman rank going down significantly. We also built a model for triple mutations as a test case despite the minimal corpus (382). This model performed well for the test set of triples, though the sparseness does not provide a strong case for true generalizability. However, this model predicts quadruple mutations better than the single or double mutant model while performing poorly on single and double mutation data. These results suggest there is some aspect of interactions captured by the FoldX energy terms that is not considered by FoldX itself as it nominally predicts additivity. One caveat is that it appears to only function for the order of mutations and one order higher (e.g. - models trained on doubles correct triple predictions, models trained on triples can enhance predictions of quadruples, etc.). This suggests that the increased complexity is partially carried over. Alternatively, it could be due to overlaps in the datasets. For example, there may be two mutations in the double mutation dataset the model learns (A+B), and those interactions are preserved in the case of triple mutations where those double mutations occur (A+B+C). Either possibility is a novel result that may advance the study and prediction of epistatic interactions.

**Figure 3:**
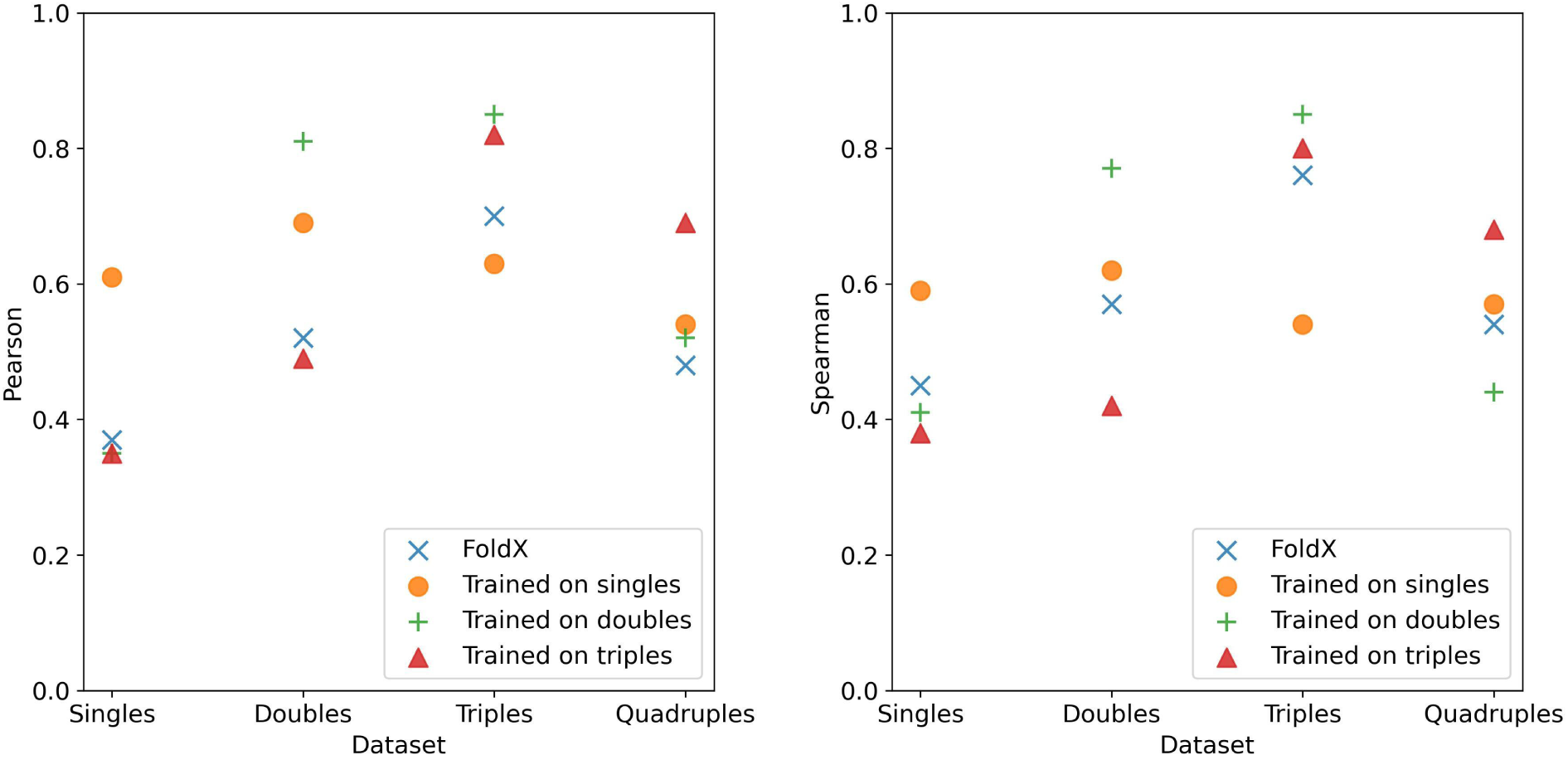
Comparison of model performance at predicting higher orders of mutations. Each model trained on N mutations (single, double, triple, etc.) was tested on the other orders of mutations to test generalizability and potential for factoring in interaction effects.

Figure 4 shows the validation results for folding and binding single mutation models and the binding double mutation model. In the case of singles, we observe generally good performance in the case of folding in terms of Pearson correlation. For Spearman rank, we see more overlap between FoldX and our corrective models. In the case of binding affinity single mutations, we observe lower performance in 3-5% of cases. Further investigation indicates these are due to one or two bad predictions in the test set causing a large deviation in the Pearson correlation. In these cases we also observe very abnormal loss performance, indicating that the model is not a good one and would be rejected outright if it were for the whole dataset or production. These subsets used the resulting hyper-parameters from the best-case model highlighting the generalizability of these tuned parameters. A full parameter search was not performed due to the cost associated.

**Figure 4:**
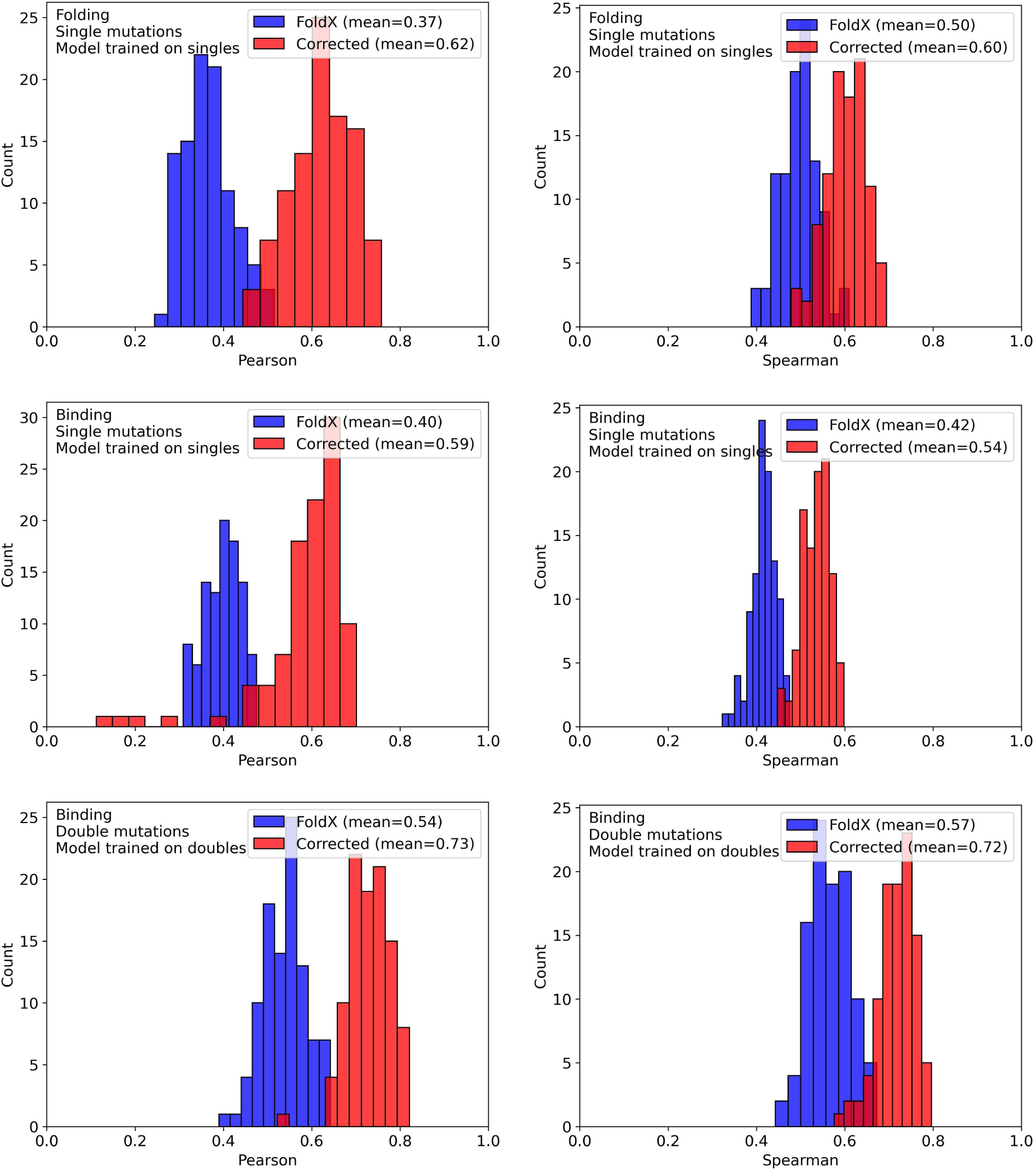
Validation results for all models tested. The x-axis is the performance metric (left: Pearson, right: Spearman) for each of 100 validation runs. The y-axis is the count. Blue and Red indicate the performance for FoldX and our model respectively.

Comparisons with other methods can be challenging due to overlapping datasets and methods curating their own subsets of SKEMPIv2 that may or may not include overlap. Several studies also include portions of their training set for evaluation inflating their results to some degree. Thankfully some methods include comparisons with external unbiased datasets. The recent method, DDAffinity, developed by Yu et al.^17^ includes comparisons with a subset of the DMS data for the Wuhan strain of SARS-CoV-2 bound to ACE2 performed by Starr et al.^40^ These are data entirely separate from SKEMPIv2 making it an excellent test case. Shown in Table 5 are our results compared to their own method and the comparisons they performed for subset S285. Of note, is that their source for the FoldX estimates, Liu et al.^39^ showed a FoldX performance of 0.39, whereas our process resulted in a Pearson of 0.46. We believe this is partially due to our methodology including iterative minimization with successive *RepairPDB* steps not performed by others and our different initial PDB prep and cleaning. In general, our model performs significantly better on this subset than any other methodology. Furthermore, the study by Yu et al. looks at higher-order mutations, however, they look at them in terms of a general subset of all multiples rather than broken down by specific order. Using our methodology on the entire SKEMPIv2 multiples dataset resulted in a performance of 0.80 compared to 0.49 with plain FoldX. Direct comparison with their methodology is difficult due to the way they broke down the SKEMPIv2 data for evaluating their model, but our predictive performance is in line with their reported results, especially when considering our validation range, meaning our methodology meets or exceeds state of the art when considering multiple mutations and generally comparable for single mutations. A significant caveat to this study, and to most similar studies, is available data. The best datasets we have accessible are SKEMPIv2.0 for binding and ProTherm4 for folding. There are other datasets discussed in the literature however these are not actually available even upon request and thus cannot be factored into studies like ours. Furthermore, the data available in these datasets is inherently biased. Datasets like SKEMPIv2.0 are compiled from formerly published experimental work. This work is usually geared towards particular systems and mutations of known significance and typically due to not include many zero-value results, or include only mutations to alanine. Ideally, we would have experimentally generated, accurate, free energy estimates comprising all possible mutations in a protein. However, the level of throughput required for an experimental method while retaining accuracy is a nontrivial task and is not frequently performed. Additionally, with such a small corpus machine learning techniques are not the most reliable. Indeed, we see more variation in our validation for double mutations than we do in singles (if one does not factor in the outliers).

**Table 5:** Results for the external SARS-CoV-2-S285 dataset for FoldX, other ML-based methods,^17^ and our corrected model. For FoldX, we included our FoldX predictions and not those provided by Liu et al.^39^

| Results for dataset S285 |  |  |  |  |
| --- | --- | --- | --- | --- |
| FoldX | RDE-Network | DiffAffinity | DDAffinity | Model Corrected |
| 0.46 | 0.403 | 0.305 | 0.456 | 0.74 |

## Conclusions

For this work, we utilized neural networks to improve the predictive capability of FoldX. Our models trained on a given order of mutations enhanced predictions for that same order of mutations, but also for higher-order mutations. Our models are able to improve predictions on an order of 50% to almost tripling predictive power. In the case of single mutations, our models improved both folding and binding affinity predictions significantly, improving correlations from Pearson correlation of 0.3 to over 0.6. For double mutations, our models did even better, with a Pearson correlation of more than 0.8. This translated to better epistasis prediction as well, almost tripling FoldX’s Pearson of 0.19 to 0.59. Using an external dataset of single mutation binding affinity with SARS-CoV-2 bound to ACE2 our methodology showed a significant performance of 0.74 compared to other machine-learning methods. Our methodology is equivalent to if not exceeding state-of-the-art with minimal added computation requirements. It can be used to great effect for building watch lists for future viruses or other diseases. Furthermore, our work highlights new aspects of epistasis and prediction not encountered in studies that use a broad multiple-mutations approach.

## Acknowledgement

Research reported in this publication was supported by the National Institute Of General Medical Sciences of the National Institutes of Health under Award Number P20GM104420. Computational resources were provided in part by Research Computing and Data Services in the Institute for Interdisciplinary Data Science at University of Idaho. The content is solely the responsibility of the authors and does not necessarily represent the official views of the National Institutes of Health.

